# MeDCycFold: A Rosetta Distillation Model to Accelerate Structure Prediction of Cyclic Peptides with Backbone N-methylation and D-amino Acids

**DOI:** 10.1101/2025.04.10.648099

**Authors:** Zhigang Cao, Tianfeng Shang, Sen Cao, Linghong Wang, Zhiguo Wang, Qingyi Mao, Jingjing Guo, Hongliang Duan

## Abstract

Cyclic peptides with backbone N-methylated amino acids(BNMeAAs) and D-amino acids(D-AAs) have gained attention for their stability, membrane permeability, and other therapeutic potentials. Currently, Rosetta can predict their structures using energy calculations, but this method is heavily time-consuming. Moreover, structural data for cyclic peptides containing BNMeAAs and D-AAs are extremely insufficient to build a data-driven structure prediction model. To address these problems, we propose MeDCycFold, a deep learning-based Rosetta distillation model by fine-tuning the AlphaFold model. First, a cyclic peptide structure dataset is constructed using Rosetta by sampling massive conformations for cyclic peptides with BNMeAAs and D-AAs and evaluating their energy scores. Then, the AlphaFold model is fine-tuned with the extended 56 BNMeAAs and D-AAs. Besides, a relative position cyclic matrix is introduced for head-to-tail cyclization in the cyclic peptides. Finally, a force field is employed to reduce clashes in the predicted structures. Empirical experiments show that our proposed MeDCycFold speeds up structure prediction by 49 times while maintaining the prediction accuracy comparable to Rosetta, which can greatly accelerate the development of cyclic peptide drugs.

## INTRODUCTION

Peptides have attracted more and more attentions in drug discovery due to their distinct properties such as high binding affinity and specificity, low toxicity, and some other therapeutic potentials^[1-5]^. However, research and development of peptide drugs are usually limited to the poor metabolic stability and membrane permeability due to their large molecular weight, high polar surface area, and some other chemical properties^[6-9]^. Compared to linear peptides, cyclic peptides are not easily to be hydrolyzed while demonstrating higher membrane permeability due to the cyclic structure. It is estimated that there are over 40 cyclic peptides or their derivatives that have been used as therapeutics^[10–13]^. To further improve the pharmaceutical properties of cyclic peptides, the most prevalent approach is to introduce various non-canonical amid acid modifications^[14]^, such as backbone N-methylation and D-AAs. Studies show that backbone N-methylation and D-AAs viable strategies to significantly improve metabolic stability, membrane permeability, and some other conformational features or properties of amide bonds ^[15-19]^.

According to Quantitative Structure-Activity Relationship (QSAR) theory, accurate structure prediction of cyclic peptides plays a crucial role during the practical cyclic peptide-based drug discovery^[20,21]^. With the rapid development of artificial intelligence, especially deep learning technologies^[22]^, many significant progresses have been made in the field of protein and peptide structure prediction^[23]^. Models such as AlphaFold ^[24,25]^, RosettaFold ^[26]^, and ESMFold^[27]^ have greatly improved the accuracy and efficiency of protein and peptide structure prediction. However, all these models are limited to protein and peptides with canonical amino acids. Recently, models like RoseTTAFold All-Atom^[28]^ have further expanded the scope of deep learning-based structural predictions to include complexes of proteins, nucleic acids, small molecules, ions, and modified residues. Building on these milestone achievements, several models for cyclic peptide structure prediction have emerged, such as AfCycDesign^[29]^ and HighFold ^[30]^, which successfully predict cyclic peptides and complexes formed by canonical amino acids with head-to-tail and disulfide bridge constraints. Subsequently, the appearance of HighFold2^[31]^ and NCPepFold^[32]^have further extended these prediction tasks to the domain of non-canonical amid acids.

However, all these AI based methods are still limited to cyclic peptides with non-canonical amid acid modifications on side chains, which are incapable of the scenario of cyclic peptides with backbone N-methylation and D-AAs. On the other side, the simple_cycpep_predict (SCP) module of Rosetta^[33,34]^ can predict cyclic peptides with BNMeAAs and D-AAs by sampling massive conformations for each cyclic peptide. Although SCP can provide relatively accurate predictions, it is heavily time-consuming due to that achieving reliable prediction precision requires thousands of structure sampling operations for average, which significantly limits the practicality of large-scale predictions.

To address these problems, we propose the MeDCycFold model (Figure 1A), which is a Rosetta distillation model^[35]^ by fine-tuning the AlphaFold framework to dramatically accelerate the structure prediction of cyclic peptides with backbone N-methylation and D-AAs with high accuracy. As shown in Figure 1B, a structural dataset of cyclic peptides with backbone N-methylation and D-AAs is constructed using the SCP module of Rosetta according to the additional dictionary of 56 BNMeAAs and D-AAs(Figure 1C and 1D). During fine-tuning the AlphaFold framework, a relative position cyclic matrix is constructed and fed into the feature embedding module to capture the cyclization information(Figure 1E). Additionally, to eliminate potential spatial conflicts in the predicted structures and minimize energy, a force field is introduced to adapt to BNMeAAs and D-AAs(Figure 1F). Compared to traditional deep learning-based structure prediction methods, MeDCycFold captures the spatial structural features of these special amino acids, filling a gap in the deep learning field, and significantly reduces structure prediction time while maintaining prediction accuracy comparable to Rosetta.

**Figure 1.**
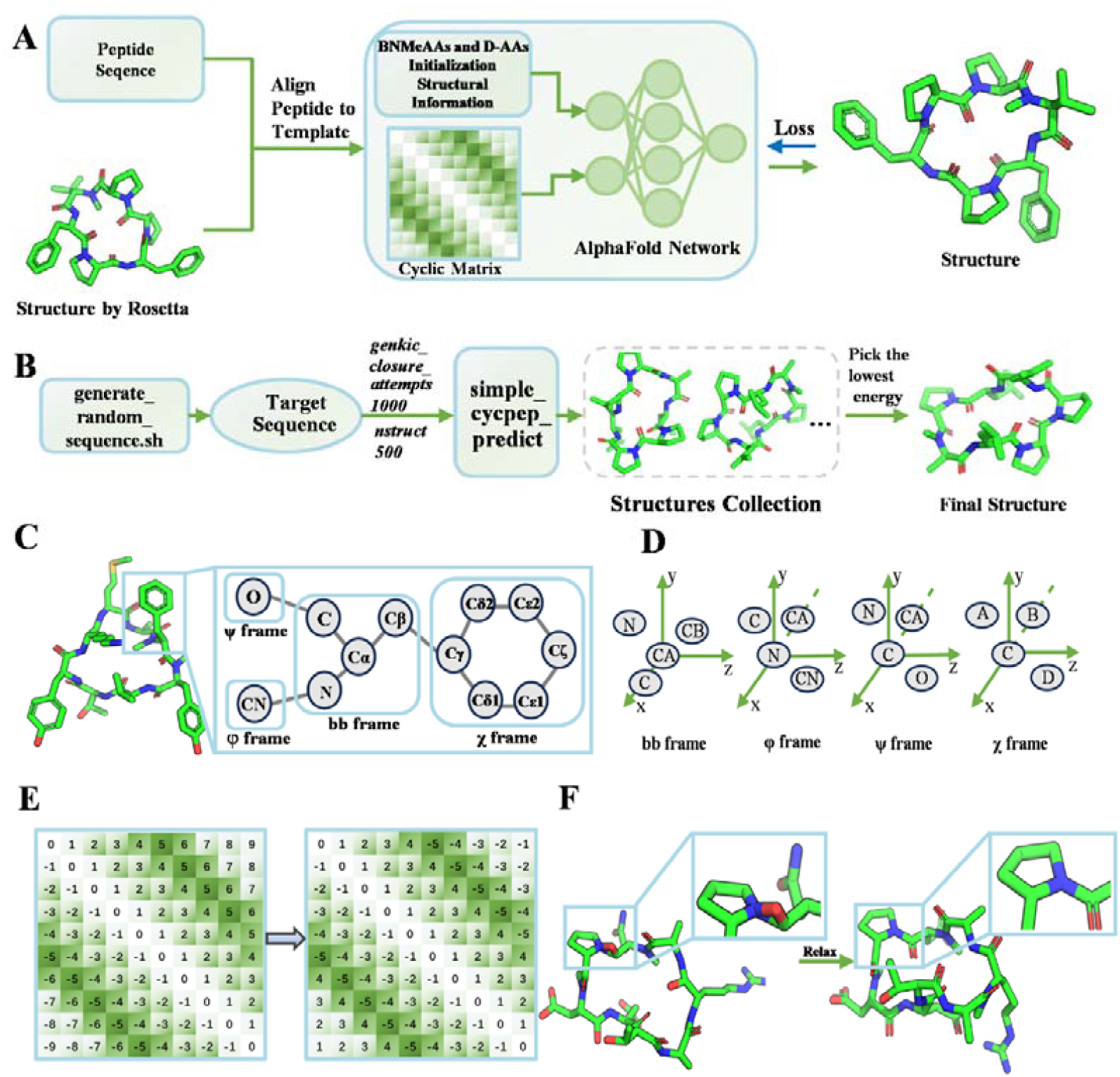
Overview of MeDCycFold. (A) Training process of MeDCycFold. (B) Workflow of dataset generation and post-processing using SCP. (C) Definition of rigid groups in BNMeAAs and D-AAs. (D) Classification of rigid groups in BNMeAAs and D-AAs. (E) Modification of relative position encoding in the prediction model to enable the prediction of cyclic peptide monomers containing BNMeAAs and D-AAs. (F) Relaxation of 3D structures containing BNMeAAs and D-AAs to eliminate spatial clashes.

## RESULTS AND DISCUSSION

A performance comparison was conducted between the MeDCycFold model and the SCP program. Subsequently, a systematic analysis was performed to examine the effects of various factors on model optimization. The impact of fine-tuning was evaluated by comparing the model’s performance before and after fine-tuning to confirm its effectiveness. The influence of incorporating a cyclic peptide coding matrix was assessed to verify its positive contribution. The effects of dataset size and the choice of loss function on model performance were analyzed to determine the optimal experimental design. Finally, the effect of force field optimization was examined. Prior to these comparative analyses, the results of the test set were directly evaluated, where the α-C RMSD reached 1.346 Å, indicating a strong agreement between the predictions and the actual structures. Furthermore, compared to the SCP program, the average prediction time was reduced from 2420.3 seconds to 48.8 seconds, highlighting the efficiency of the model.

### Performance comparison between MeDCycFold and SCP

Since the dataset was generated under the parameters *nstruct = 500* and *cyclic_peptide:genkic_closure_attempts = 1000*, a comparison was conducted between the prediction accuracy and computational time of SCP under these conditions and those of MeDCycFold. It is important to note that Rosetta’s prediction accuracy is highly dependent on parameter settings. The chosen parameters were determined based on a balance between computation time and available resources, meaning the results presented here do not represent the best possible accuracy of SCP.

As shown in Figures 2A and 2B, the average α-C RMSD of MeDCycFold is 0.204 Å higher than that of SCP, while the average computation time is reduced by 2371.5 seconds. Although SCP produces slightly lower RMSD values for different structures, MeDCycFold demonstrates significantly higher efficiency, making it more suitable for large-scale predictions while maintaining competitive accuracy. Notably, for 12 structures, including D9.16, D8.26, D10.1, D8.17, D8.19, D9.24, D11.25, D8.25, D8.6, D8.5, D8.21, and D8.15, more accurate α-C predictions are achieved compared to SCP. In addition, the median α-C RMSD of MeDCycFold is 1.098 Å, which is lower than the 1.119 Å obtained by SCP (Table S1). It is worth noting that 13 of the 28 test cases, the structures predicted by MeDCycFold achieved α-C RMSD below 1Å when compared to the native structure. To further illustrate performance differences, several structures were visualized. Figures 2C, 2D, and 2E present examples where MeDCycFold consistently outperforms SCP, achieving lower RMSDs in all evaluated metrics, including α-C, backbone, and all-atom predictions (left: MeDCycFold; right: SCP; same applies to Figures 2F, 2G, and 2H). Notably, for D8.26, the α-C RMSD improved from 2.073 Å to 0.990 Å. Additionally, Figures 2F, 2G, and 2H highlight cases where MeDCycFold demonstrates superior accuracy in predicting α-C atoms.

**Figure 2.**
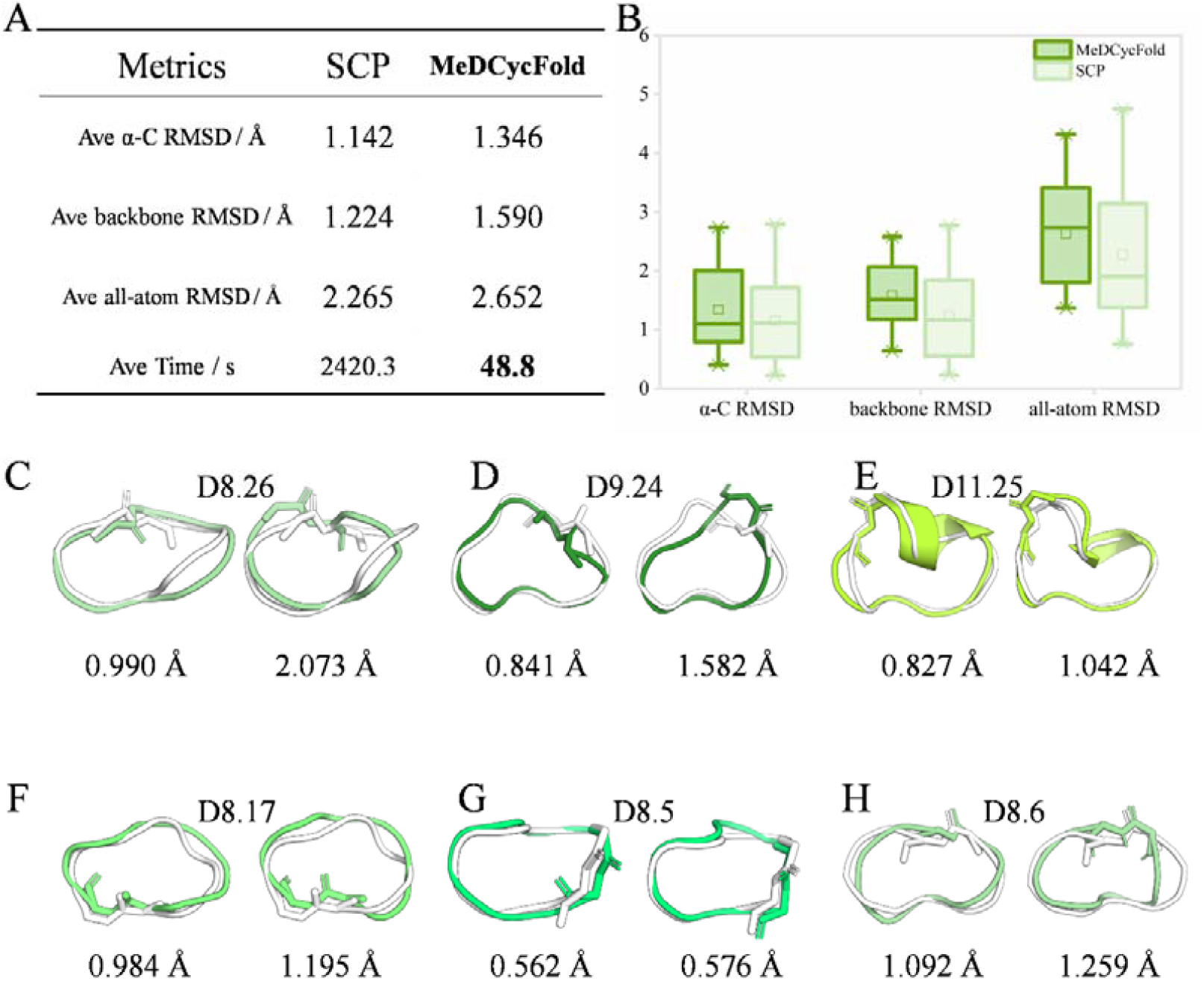
(A) Comparison of RMSD and computational time between MeDCycFold and SCP. (B) RMSD distribution of α-C, backbone, and all-atom measurements for both methods. (C–H) Structural alignment of selected cyclic peptides, where the native structures are shown in gray, MeDCycFold predictions on the left, and SCP predictions on the right. The displayed examples include (C) D8.26, (D) D9.24, (E) D11.25, (F) D8.17, (G) D8.5, and (H) D8.6, with their corresponding RMSD provided.

### Comparison before and after fine-tuning

Figure 3A and 3B show the results of the RMSD comparison. The fine-tuned model performs best with an average α-C RMSD of 1.346 Å, an average backbone RMSD of 1.590 Å, and an average all-atom RMSD of 2.652 Å. The median α-C RMSD is 1.098 Å (Table S1). In contrast, the model without fine-tuning had an average α-C RMSD of 2.218 Å, an average backbone RMSD of 2.253 Å, and an average all-atom RMSD of 3.692 Å. The precision of each was, respectively, improved by improved by 0.872 Å, 0.663 Å, and 1.04 Å. 25 structures were predicted more accurately after fine-tuning compared to those before fine-tuning in the 28 test sets, demonstrating a huge improvement in the performance of the model after fine-tuning. As shown in Figure 3C–H, several fine-tuned predicted structures achieve an α-C RMSD of less than 1 Å, including D7.6 (0.438 Å) and D8.1 (0.400 Å), demonstrating improved structural accuracy compared to the model without fine-tuning.

**Figure 3.**
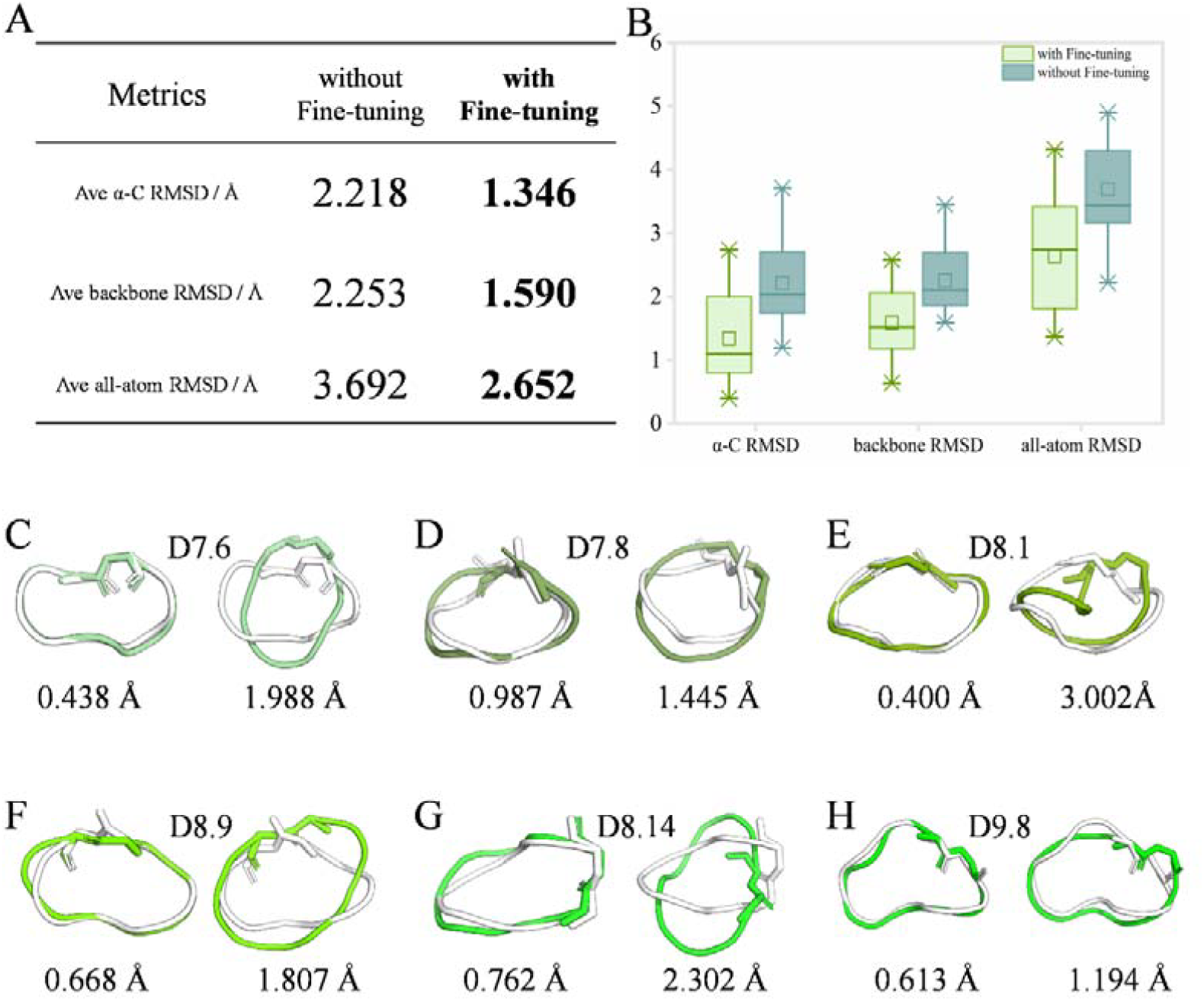
(A) Comparison of RMSD between the model with fine-tuning and the model without fine-tuning. (B) RMSD distributions for α-C, backbone, and all-atom measurements, highlighting the enhancement in structural accuracy achieved through fine-tuning with the cyclic peptide matrix. (C–H) Structural comparisons of selected cyclic peptides, where native structures are shown in gray, predictions generated by the model with fine-tuning are displayed on the left, and predictions from the model without fine-tuning are shown on the right. The selected examples include (C) D7.6, (D) D7.8, (E) D8.1, (F) D8.9, (G) D8.14, and (H) D9.8, with their respective RMSD values presented below each structure.

### Analysis of the role of the relative position cyclic matrix

When testing cyclic peptides, the initial AlphaFold model generates linear peptide structures, making it unsuitable for generalizing to cyclic peptides with non-canonical modifications. This limitation is reflected in the high average α-C RMSD of 5.819 Å across the entire test set. However, the incorporation of the cyclic peptide coding matrix reduces the α-C RMSD to 2.218 Å, demonstrating its significant impact. Figures 4A and 4B present the complete RMSD distribution.

**Figure 4.**
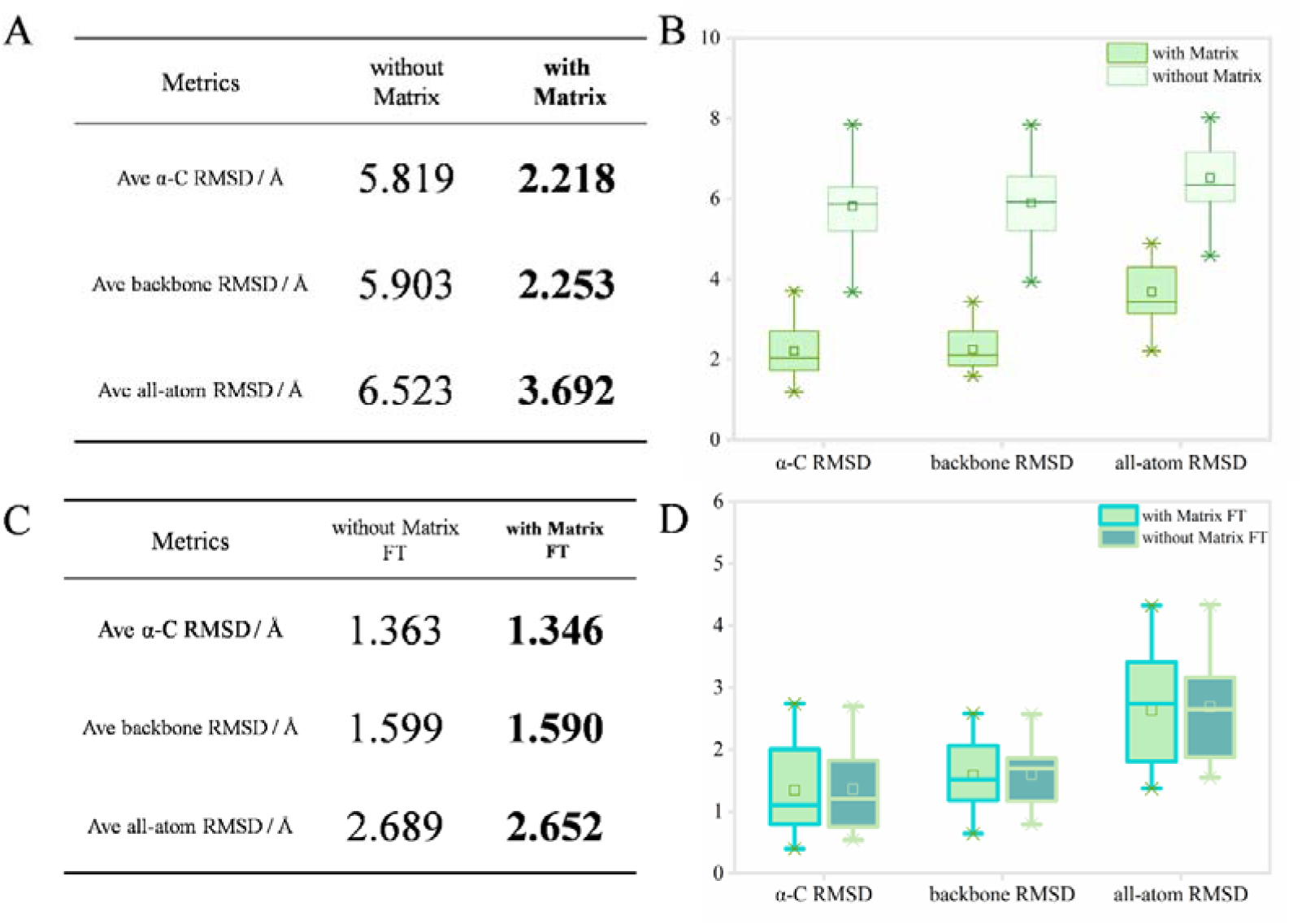
(A)Comparison of RMSD results with and without matrix testing. (B) Distribution of RMSD with and without matrix testing.(C) Comparison of RMSD results with and without matrix fine-tuning. (D) Distribution of RMSD with and without matrix fine-tuning

During the fine-tuning phase, models with and without the cyclic peptide coding matrix were trained using the same dataset, and their performance was evaluated on the test set. As shown in Figures 4C and 4D, the α-C RMSD of the model fine-tuned with the cyclic peptide coding matrix was 1.346 Å, whereas the model fine-tuned without it reached 1.363 Å. The RMSD for the model with cyclic peptide matrix fine-tuning were slightly lower than those of the model without the matrix fine-tuning: the RMSD for the backbone was 1.590 Å compared to 1.599 Å, and the RMSD for all atoms was 2.652 Å compared to 2.689 Å.

### Impact of dataset size on model performance

To investigate the impact of dataset size on model performance, the prediction accuracy of the model was evaluated across various training dataset scales, using α-C RMSD as the performance metric. Figure 5A illustrates the relationship between training dataset size and α-C RMSD, with the horizontal axis representing dataset sizes (100, 500, 1000 and 2500) and the vertical axis representing α-C RMSD. The specific data points are as follows: at a dataset size of 100, the α-C RMSD is 1.927 Å; At 500, it is 1.797 Å; At 1000, it is 1.544 Å; And at 2500, it reaches 1.346 Å.

**Figure 5.**
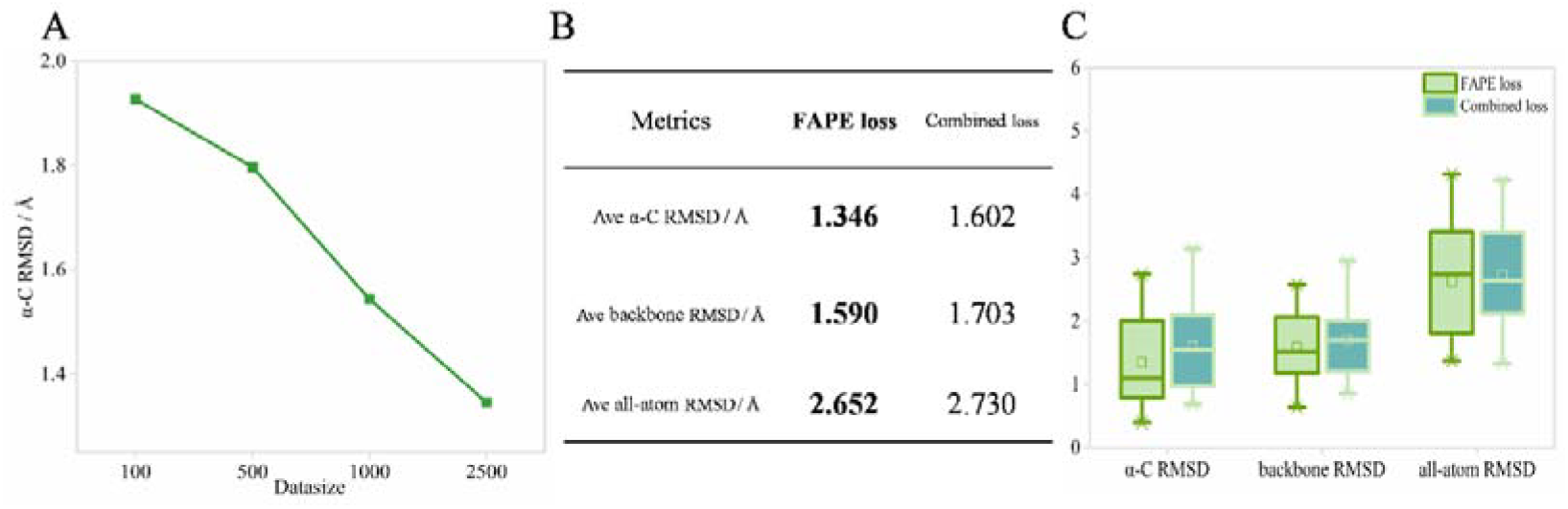
(A)Changes in α-C RMSD in different data set sizes.(B)Comparison of RMSD using FAPE loss and combined loss. (C) Distribution of RMSD using FAPE loss and combined loss.

As shown in Figure 5A, α-C RMSD exhibits a nonlinear decreasing trend with increasing dataset size, indicating a progressive improvement in model performance. Specifically, when the dataset size increases from 100 to 500, α-C RMSD decreases from 1.927 Å to 1.797 Å, a reduction of 0.13 Å. This suggests that the increase in data volume contributes to some improvement in model performance. However, due to the limited data size, the model may not yet fully capture the complex features of cyclic peptide structures, leading to relatively high prediction errors. As the dataset size further increases to 1000, α-C RMSD drops to 1.544 Å, a more significant reduction of 0.253 Å, demonstrating that the additional data allows the model to better capture structural features. Finally, when the dataset size increases from 1000 to 2500, α-C RMSD further decreases to 1.346 Å, a reduction of 0.198 Å, indicating a continued improvement in model performance and reaching its optimal performance at this point. This suggests that at a dataset size of 2500 samples, the model can adequately learn the diversity of cyclic peptide structures while avoiding the risk of overfitting. Overall, α-C RMSD decreases from 1.927 Å (Dataset size = 100) to 1.346 Å (Dataset size = 2500), a total reduction of 0.581 Å, demonstrating that the model’s prediction accuracy significantly improves as the dataset size increases.

### Performance comparison of loss functions in fine-tuning for structures containing BNMeAAs and D-AAs

Two fine-tuning strategies were developed to enhance model performance. The first strategy utilizes the multi-objective loss function inherent to AlphaFold, referred to as the combined loss, incorporating the Frame-Aligned Point Error (FAPE) and several auxiliary losses. The second strategy employs FAPE exclusively as a single loss function for fine-tuning. Both strategies were tested across different dataset sizes (100, 500, 1000, and 2500), and the fine-tuned models were evaluated on a test set comprising 28 real cyclic peptide structures, with performance assessed using the average RMSD as the metric. Optimal performance was achieved with both strategies at a dataset size of 2500. Figures 5B and 5C present the RMSD comparison and RMSD distribution, respectively, for a dataset size of 2500. The single FAPE loss strategy outperformed the combined loss strategy, achieving an average α-C RMSD of 1.346 Å (versus 1.602 Å), a backbone RMSD of 1.590 Å (versus 1.703 Å), and an all-atom RMSD of 2.652 Å (versus 2.730 Å).

The outperformance of the single FAPE loss strategy over the combined loss strategy can be attributed to the complexity of BNMeAAs and D-AAs, the constraints imposed by dataset size, and the optimization objectives of the loss function. The inclusion of 56 BNMeAAs and D-AAs substantially increases the diversity of the input space, while the auxiliary losses in the combined loss function may fail to effectively adapt to the sequence and structural characteristics of BNMeAAs and D-AAs, potentially causing interference during training. In contrast, the single FAPE loss focuses on structural alignment errors, directly optimizing the geometric consistency between predicted and actual structures. This approach may be better suited for the modeling requirements of cyclic peptides and BNMeAAs and D-AAs in the present context.

### Relaxation of structures containing BNMeAAs and D-AAs

AlphaFold is a groundbreaking deep learning model capable of accurately predicting three-dimensional structures of biomacromolecules. For the initially generated structures, AlphaFold typically employs an Amber force field-based relaxation process (Amber-relax) to perform energy minimization, enhancing the stereochemical quality of the predicted structures. This relaxation step eliminates atomic clashes, optimizes bond lengths and angles, and refines side-chain conformations, ensuring the physical and chemical plausibility of the structures. However, the standard Amber-relax workflow is limited to the 20 canonical amino acids due to the availability of predefined force field parameters. When structures contain BNMeAAs and D-AAs, the absence of corresponding parameters prevents the standard Amber-relax process from proceeding, thereby restricting further refinement of protein structures with chemical modifications or BNMeAAs and D-AAs included. This limitation poses a challenge to the accurate prediction and design of proteins containing BNMeAAs and D-AAs.

To address this limitation, an enhanced Amber-relax strategy was developed by integrating quantum chemical calculations with molecular mechanics. Specifically, Gaussian software and AmberTools were employed to derive Amber-compatible parameter files. Subsequently, the tleap module in AmberTools was used to generate topology and coordinate files for the target structures. Finally, energy minimization of the full structure was performed using the efficient molecular simulation engine OpenMM. Through these steps, force field parameters for 56 BNMeAAs and D-AAs were successfully integrated into the Amber-relax workflow, enabling effective processing and optimization of predicted structures containing BNMeAAs and D-AAs.

As shown in Figure 6A, the relaxed structures showed significantly improved quality relative to the initial unrelaxed ones. Specifically, the enhanced Amber-relax protocol effectively reduced atomic clashes within the structures (Figure 6B) and optimized side-chain conformations through energy minimization. Detailed results are provided in Supplementary Materials (Table S2).

**Figure 6.**
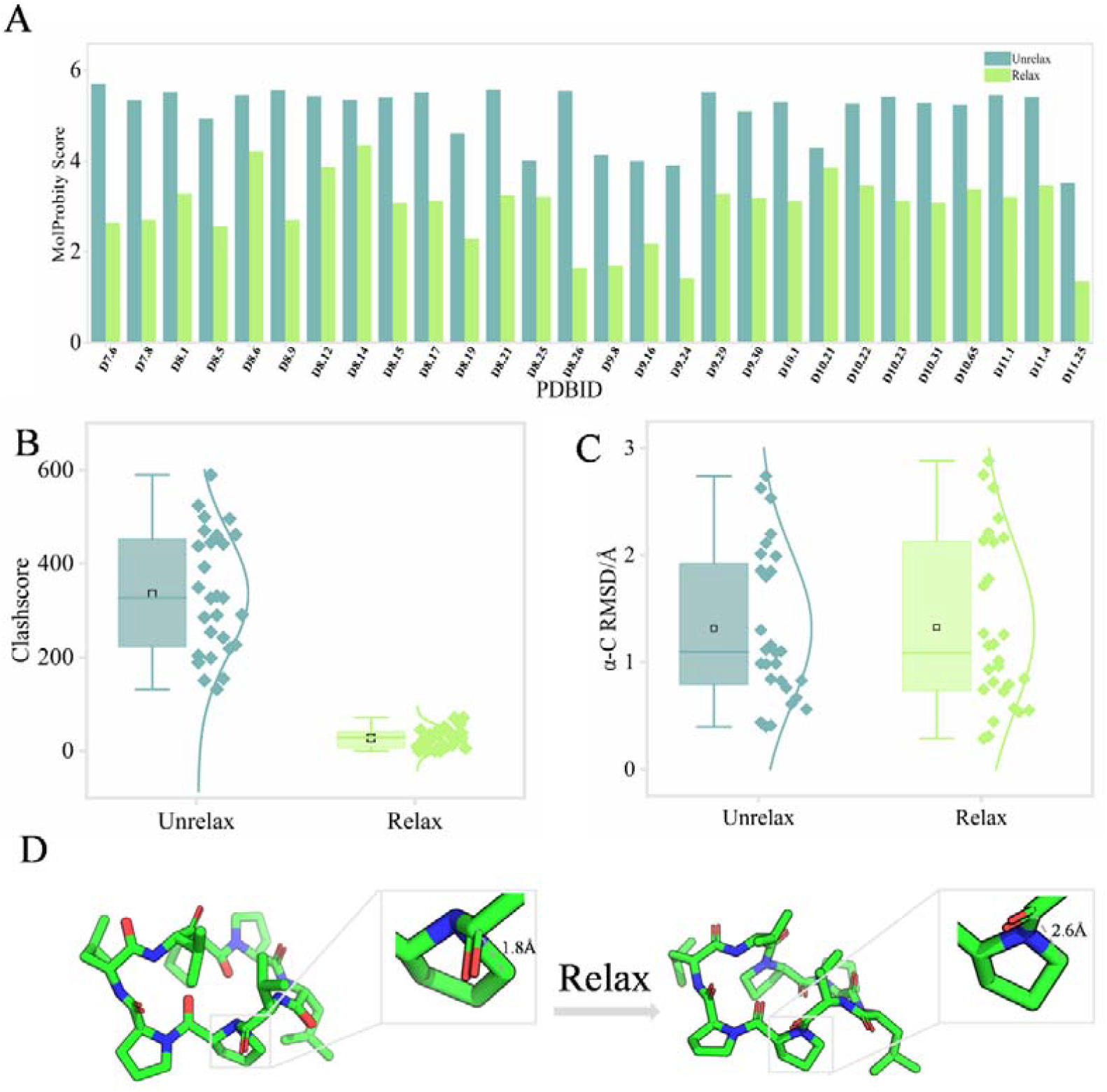
(A) Bar chart of MolProbity scores before and after relaxation on the test set.(B) Distribution of clashscore before and after relaxation on the test set.(C) Distribution of α-C RMSD before and after relaxation on the test set.(D) The distance between the C atom of PRO at position 8 and the CD atom of the residue at position 3 in D8.12 was relaxed from 1.8 Å to 2.6 Å after the relaxation process.

**Figure 7.**
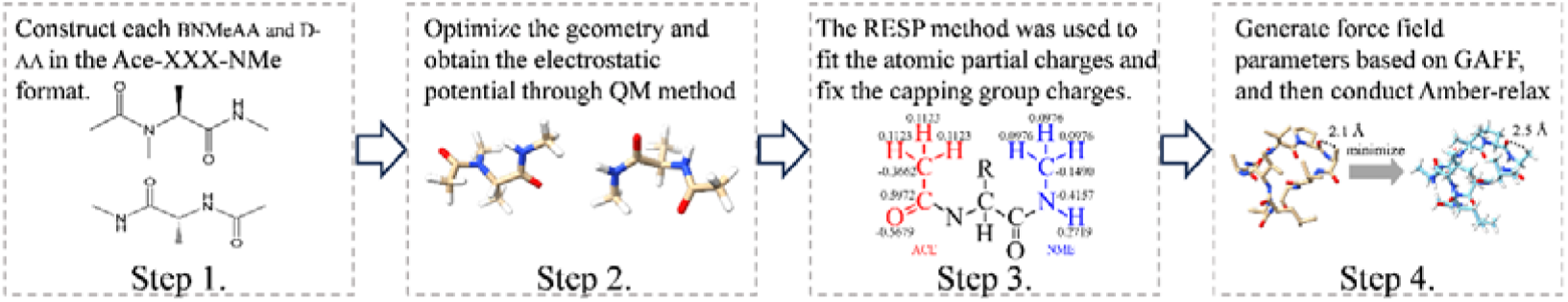
Preparation of charges and parameters for BNMeAAs and D-AAs.

To assess the effect of Amber-relax on prediction accuracy, we compared the predicted structures before and after relaxation with the corresponding native structures. As illustrated in Figure 6C, Amber-relax basically did not change the overall accuracy of the predicted models, as indicated by the average α-C RMSD value of 1.35 Å for the unrelaxed and relaxed structures, more detailed results were provided in Supplementary Materials (Table S3). This result indicates that the primary effect of Amber-relax was the improvement of local geometric features, such as side-chain positioning and stereochemical plausibility, without significantly altering the overall structural prediction accuracy, consistent with previous findings.

Furthermore, detailed local interactions of typical side-chain atoms were examined (Figure 6D). In the unrelaxed models, clearly unreasonable nonbonded interactions between side-chain atoms were observed. These local structural issues were effectively resolved following the Amber-relax optimization procedure. Collectively, our enhanced Amber-relax workflow significantly improved both the reliability and local geometric accuracy of the predicted structures.

By combining QM calculations with deep learning, our work substantially extends AlphaFold’s applicability to structures containing BNMeAAs and D-AAs, offering new possibilities for protein design and structural optimization. However, it was observed that some relaxed structures retained minor steric clashes or geometric irregularities, suggesting that future studies could incorporate more refined optimization techniques, such as extended molecular dynamics simulations, to further improve the structural reliability and physical realism of the predicted structures.

## METHODS

### Experiment settings

The MeDCycFold model and Rosetta with version 3.31 are running on Ubuntu 20.04.4 LTS. The hardware setup includes an Intel i9-12900KF CPU (24 cores), an RTX 4090 GPU (24GB), 192GB of RAM, and a 12TB hard drive.

### Datasets

In this study, we utilized the SCP to generate the model training and validation datasets, as shown in Figure 1B. First, we constructed a random sequence generation script: generating a list of 39 amino acids without methylation and a list of 37 amino acids with methylation.Amino acids were then randomly selected from these lists to obtain random sequences. To ensure the rationality and diversity of the generated cyclic peptide structures while considering computational resources, we set key parameters in the SCP simulation script: *nstruct = 500* and *cyclic_peptide:genkic_closure_attempts = 1000*. Subsequently, structure prediction was performed, and the lowest-energy structures were selected. Since SCP assigns the same names to BNMeAAs and their unmethylated precursors, we modified the obtained PDB files to ensure consistency with the BNMeAAs and D-AAs dictionary in our model. Ultimately, 2750 cyclic peptide structures were randomly generated. From these, 2500 structures were randomly selected as the training set, 250 as the validation set, and the test set comprised 28 X-ray crystal structures provided in the study by David Baker (2022)^[36]^. Specifically, we selected 28 cyclic peptides with BNMeAAs and D-AAs that matched the 21 types included in our dataset to validate the accuracy and reliability of our proposed method in real-world structural predictions.

### Model Architecture

MeDCycFold is based on AlphaFold and fine-tuned with its parameters. To predict cyclic peptides containing BNMeAAs and D-AAs, we extended AlphaFold by incorporating an dictionary of BNMeAAs and D-AAs, adding a cyclic peptide matrix, and modifying the loss function.

#### (a) Addition of a dictionary of BNMeAAs and D-AAs

We expanded AlphaFold’s amino acid dictionary by introducing structural information for BNMeAAs and D-AAs. Specifically, 19 BNMeAAs were obtained by methylating the nitrogen atom of canonical amino acids(excluding PRO);19 D-AAs were constructed by generating enantiomers of canonical amino acids(excluding GLY); And 18 D-BNMeAAs were obtained by methylating the main-chain nitrogen of D-AAs (excluding D-GLY). Ultimately, we added 56 BNMeAAs and D-AAs, enabling the model to handle these complex molecules. The names and code symbols of these 56 amino acids are provided in Table S4. The structural module of AlphaFold uses single and pairwise representations generated by the Evoformer module to predict the torsion angles of amino acid residues. These predicted torsion angles, along with predefined rigid groups and atomic coordinates, help the model accurately construct atomic positions in three-dimensional space, simplifying structural prediction complexity. To extend the model for BNMeAAs and D-AAs, we defined the rigid groups of these residues and initialized their atomic coordinates.

Atoms in amino acids are classified into five rigid groups based on their dependence on specific torsion angles (Table S5). The backbone rigid groups (bb framework), which consist of Cα, Cβ, C, and N atoms, serve as the structural core. The φ group, which was initially used to compute the coordinates of the hydrogen atom on the amine group, has been repurposed to determine the position of the CN methyl group due to the substitution of hydrogen. The ω group, which is associated with the hydrogen atom on Cα, is excluded, as hydrogen atoms cannot be predicted by the model. The ψ rigid group, which includes the oxygen atom of the carboxyl group, is distinct from the χ rigid group, which comprises all side-chain atoms (Figure 1C). The initial coordinates of these rigid groups, which are derived from the structure predicted by SCP, provide a foundation for further structural refinement.

As illustrated in Figure 1D, the orientation of the bb framework, which is centered on α-C, positions C along the positive x-axis while constraining N to the xy-plane, thereby allowing the coordinates of Cβ to be determined. The φ group, which is referenced to N as the origin, is defined by placing Cα along the negative x-axis and constraining the adjacent C atom to the xy-plane, thereby determining the position of CN. The ψ rigid group, which is initialized with respect to C, is defined by positioning Cα along the negative x-axis and constraining the nitrogen atom to the xy-plane, thereby determining the oxygen atom’s position. The side-chain χ rigid group, which is subdivided into four subgroups based on distinct torsion angles, accommodates amino acids with more than four torsion angles by neglecting those with negligible influence. Within each subgroup, the third atom is set as the origin, the second atom is aligned along the negative x-axis, and the first atom is positioned within the xy-plane. The relative position of the fourth atom is rotated into the xy-plane using the rotation matrix *Rx*, as described below:

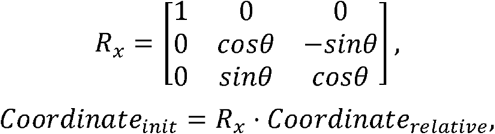

where θ represents the rotation angle around the x-axis to align the fourth atom with the xy-plane.

#### (b) Introduction of relative position cyclic matrix

To accommodate the unique structural characteristics of cyclic peptides, we incorporated a cyclic matrix into the AlphaFold model, inspired by AfCycDesign^[29]^. This modification enables the model to explicitly capture the cyclic constraints of cyclic peptides, enhancing its ability to predict such structures. In the original AlphaFold model, relative positional encoding defines sequence distances between amino acids. For linear peptides and cyclic peptides, the sequence distance between neighboring residues is 1, while the distance between N-terminal and C-terminal residues is the sequence length - 1. For instance, in the linear peptide sequence ACDEFGHIJK, the sequence distances between residue A and CDEFGHIJK are 1, 2, 3, 4, 5, 6, 7, 8, and 9. However, this encoding fails to capture the cyclic nature of peptides, leading to suboptimal performance. To address this issue, we designed a custom *N×N* cyclic offset matrix that introduces cyclic information into the relative positional encoding. In this matrix, the sequence distance between the first and last residues of a cyclic peptide of length N is redefined as 1, simulating the peptide’s closure. For instance, after applying this encoding, the sequence distances between residue A and CDEFGHIJK in a cyclic peptide are redefined as 1, 2, 3, 4, -5, -4, -3, -2, and -1, with negative signs indicating opposite directionality (Figure 1E). This encoding correctly represents the relationship between terminal residues and comprehensively captures the cyclic nature of peptides. The cyclic offset matrix undergoes one-hot encoding and linear projection before being embedded into AlphaFold’s Evoformer module as part of the pairwise features. This adjustment significantly improves the model’s ability to capture global cyclic peptide characteristics.

#### (c) Loss function

AlphaFold is trained in an end-to-end manner, with gradients derived from the Frame Aligned Point Error (FAPE) ^[24]^loss and several auxiliary losses, forming combined loss that includes the FAPE loss, the auxiliary loss from the Structure Modul, the distogram prediction loss, the masked MSA prediction loss, and the model confidence loss. Through experimentation, we opted to use only the FAPE loss in our final model.

### Relaxation

#### (a) BNMeAAs and D-AAs force field

Force field parameters for BNMeAAs and D-AAs were developed following the Amber protocol (https://carlosramosg.com/amber-custom-residue-parameterization). Initially, each BNMeAA and D-AA was built in GaussView6 using the Ace-XXX-NMe capping approach (Step 1of Figure7)^[37]^. Geometry optimization was performed in the gas phase at the B3LYP/6-31G(d) level, followed by electrostatic potential (ESP) calculations via single-point Hartree–Fock (HF) computations at the 6-31G(d) basis set (Step 2 of Figure7)^[38]^, aligning with the standard Amber methodology for deriving partial atomic charges. The ESP was extracted from Gaussian output files using the espgen module in AmberTools^[39]^, and atomic partial charges were fitted using the restrained electrostatic potential (RESP) procedure, with charges of the ACE and NME capping groups constrained to match standard Amber force field values (Step 3 of Figure7). Finally, capping groups were removed using the prepgen module in AmberTools, and topology files (.prepin and .frcmod) were generated based on the generation of General Amber Force Field (GAFF), ensuring atom naming consistency with corresponding PDB files (Step 4 of Figure7).

#### (b) Energy minimization

The predicted peptide structures were loaded into the tleap module of AmberTools, where the Amber ff14SB force field, GAFF, and the newly developed parameters specific to BNMeAAs and D-AAs were integrated to generate the topology and coordinate files required for energy minimization^[40]^.

Energy minimization was performed using OpenMM^[41]^ (version 7.7.0) to relax the predicted structures while preserving their initial conformations as closely as possible. The simulation system was constructed using the AmberPrmtopFile and AmberInpcrdFile classes in OpenMM, with nonbonded interactions set to NoCutoff and all bonds involving hydrogen atoms constrained via HBonds. To prevent significant structural deviations during minimization, harmonic positional restraints were applied to backbone atoms. The minimization process continued until convergence, with an energy tolerance of 2.39 kcal/mol.

### Metrics

#### (a) RMSD

In this study, we use α-C RMSD, backbone RMSD and all-atom RMSD as the evaluation metrics to quantify the deviation of the model predicted structure from the reference structure. RMSD calculation are conducted in three approaches, and formulated as follows:

*N* denotes the numbers of all atoms, backbone atoms (C, CA, N, O and CN) and α-C atoms for three approaches respectively.

#### (b) Structure assessments

Independent structural assessments were conducted using the MolProbity online server (^https://molprobity.biochem.duke.edu/)[40]^. It evaluates the stereochemical quality and accuracy of macromolecular structures by analyzing key parameters, including atomic contacts, molecular geometry, and backbone torsion angles. MolProbity is widely used in structural biology and provides quantitative and reliable evaluations of biomacromolecule structures.

## CONCLUSION

Cyclic peptides containing BNMeAAs and D-AAs exhibit enhanced membrane permeability and stability, making them promising candidates for drug development. Structural elucidation of these peptides not only aids in understanding their interactions with protein targets but also provides a critical foundation for novel drug design. However, the scarcity of experimentally determined structures for such peptides has significantly hindered the application of deep learning models in this domain.

In this study, the Rosetta SCP program was utilized to generate a dataset of cyclic peptides containing BNMeAAs and D-AAs. The AlphaFold model was subsequently fine-tuned on this dataset, enabling it to predict these cyclic peptide structures with higher accuracy. This work fills a gap in deep learning-based predictions of cyclic peptides with BNMeAAs and D-AAs and highlights the potential of the model framework for broader applications involving other noncanonical peptide structures. Additionally, the study provides a novel strategy for generating computational datasets to train specialized deep learning models.

Despite these advancements, the model has certain limitations. Specifically, while it improves prediction efficiency, it has yet to surpass Rosetta’s SCP in accuracy. This is likely due to the reliance on Rosetta-generated data for fine-tuning, which may constrain model performance. The analysis suggests that training on higher-quality datasets generated with more refined Rosetta parameters could enhance prediction accuracy. Furthermore, computational resource limitations may have impacted model performance, and future improvements in computational infrastructure could further optimize both data generation and model fine-tuning.

Importantly, the model establishes a scalable framework that integrates computational tools with deep learning, extending beyond cyclic peptides containing BNMeAAs and D-AAs. This approach reduces the dependency on experimental structural data and alleviates the resource-intensive nature of experimental structure determination, thereby facilitating the study and development of peptide-based therapeutics.

In conclusion, this study demonstrates the substantial potential of combining computational tools with deep learning to address complex cyclic peptide structure prediction challenges, providing valuable insights and guidance for future research on non-canonical amid acid peptides.

## Author contributions

Z.C., T.S., and H.D. conceptualized ideas, proposed methods, and wrote the manuscript. Z.C., T.S., and S.C. investigated and implemented the deep learning programs. L.W., Z.W., Q.M., and J.G. completed the data collection. All authors have read and approved the final manuscript.

## DATA AVAILABILITY

The MeDCycFold code can be found at https://github.com/hongliangduan/MeDCycFold.

## SUPPLEMENTARY MATERIALS

Tables S1 to S9

